# Deep Learning Identifies BMI-Associated Morphologic Features in Breast Cancer Tissue

**DOI:** 10.64898/2026.11.10.765089

**Authors:** Edwin A. Heredia, Lara Papasian, John Paine, Shiva Patre, Thomas J. Jönsson, Thomas Keller, Zihang Fang, Valerie H. Narumi, Irika Katiyar, Ian T. Lagerstrom, Jamie Lombardo, Brianna Simental, Brian Tran, Denisse Sibrian, Olga Castellanos, Justin Wells, Jerry S. H. Lee, Craig Shriver, David B. Agus, Reva K. Basho, Naim Matasci

## Abstract

Multiple epidemiological studies have established obesity as an adverse factor in breast cancer incidence, progression, and clinical outcomes. However, the biological mechanisms underlying these associations remain incompletely understood. In peritumoral adipose tissue, obesity is associated with chronic inflammation, macrophage recruitment, and broader alterations in the local microenvironment. Whether these obesity-associated changes are accompanied by distinct morphologic features within the tumor and non-adipose stromal compartments remains unclear. In this study, we investigated whether an AI-based pathology foundation model could support the development of machine learning algorithms capable of identifying obesity-associated morphologic patterns in histopathology images, with a particular focus on tumor and stromal regions. In a cohort of 195 patients with breast cancer, machine learning models using vector embeddings from non-adipose stromal regions classified high BMI (≥30 kg/m^2^) versus low BMI (<25 kg/m^2^) with an average AUC of 0.78 (95% CI: 0.73–0.83), consistently outperforming models based on tumor-only or combined tumor–stroma regions. These obesity-associated stromal signatures were observed across subgroups defined by hormone-receptor status and age, indicating that they are not restricted to specific clinical populations. Complementary embedding-based dissimilarity analysis further showed that higher BMI was associated with reduced inter-patient heterogeneity in both tumor and stromal tissues. Overall, these findings suggest that obesity is associated with distinct morphologic alterations in both tumor and non-adipose stromal tissues. Notably, the stromal compartment emerged as a particularly informative indicator of obesity-associated morphology and demonstrated reduced inter-patient heterogeneity among individuals with high BMI.

## 1 Introduction

Obesity is a major public health concern in the United States and other developed countries, affecting approximately 40% of the US adult population (Emmerich, et al., 2024). People with obesity are at increased risk for numerous serious conditions, including cardiovascular disease, type 2 diabetes, osteoarthritis, and certain cancers. (Dixon, 2010). The body mass index (BMI), defined as a person’s weight in kilograms divided by the square of their height in meters (kg/m^2^) provides a measure of obesity, and despite some known limitations (Romero-Corral, et al., 2008), is widely used to screen for obesity in clinical practice. The BMI scale categorizes individuals into four groups: underweight (BMI < 18.5), healthy weight (18.5 ≤ BMI < 25), overweight (25 ≤ BMI < 30), and obese (BMI ≥ 30).

High BMI is consistently associated with an increased risk of postmenopausal breast cancer, particularly estrogen receptor (ER)-positive disease (Reeves, et al., 2007; Munsell, et al., 2014). In contrast, obesity appears protective against most premenopausal breast cancers, although it may increase the risk of triple-negative breast cancer (TNBC) (Munsell, et al., 2014; Pierobon & Frankenfeld, 2013; García-Estévez, et al., 2021). Obesity is also linked to poorer prognosis in early breast cancer characterized by human epidermal growth factor receptor 2 (HER2) positivity (Martel, et al., 2021) and with reduced treatment efficacy (Lee et al., 2019).

Despite strong epidemiological evidence, the mechanisms linking obesity to breast cancer remain incompletely understood (Quail & Dannenberg, 2019; Kolb & Zhang, 2020). In obesity, hypertrophic adipocytes undergo stress and necrosis, recruiting CD68+ macrophages that form crown-like structures (CLS), a hallmark of chronic adipose inflammation. CLS are associated with increased aromatase activity, pro-inflammatory cytokines (TNF-α, IL-6), and altered adipocyte–tumor crosstalk, collectively promoting breast cancer progression (Kolb & Zhang, 2020; Lyu, et al., 2022; Kothari, et al., 2020; Morris, et al., 2011).

Several studies have identified obesity-associated changes in the breast tumor microenvironment, particularly within peritumoral adipose tissue. Histological analyses consistently show that crown-like structures (CLS), adipocyte size, and macrophage infiltration are positively associated with BMI (Morris, et al., 2011; Iyengar, et al., 2015; Sun, et al., 2012; Chang, et al., 2021; Maliniak, et al., 2021; Faria, et al., 2020). At the molecular level, obesity is also associated with enrichment of inflammatory and metabolic pathways, supporting a link between obesity and altered tumor microenvironment (TME) biology (Sun, et al., 2012).

While most of the studies of obesity-related markers in the TME have focused predominantly on peritumoral adipose tissue, it is not known if obesity also induces observable changes in the tumor and associated non-adipose stroma. In this study, we hypothesized that obesity-associated morphologic features are detectable in tumor and non-adipose stromal tissue using modern artificial intelligence (AI) models trained to identify histologic patterns predictive of BMI status. We further hypothesized that such features may be more pronounced in postmenopausal women and in patients with breast cancers characterized by estrogen receptor (ER) positivity and HER2 negativity (ER+/HER2−).

Recent advances in AI-based computational pathology have enabled foundation models that learn generalizable morphologic representations from unlabeled histopathology images (Ochi, et al., 2025; Li, et al., 2025; Bilal, et al., 2025). In this study, we used the CanvOI pathology foundation model (Zalach, et al., 2024) to generate vector representations from WSIs obtained from 195 breast cancer patients. These representations were further analyzed using conventional machine learning classifiers to identify associations between tissue morphology and BMI, categorized as low (<25 kg/m^2^) or high (≥30 kg/m^2^), across tumor and stromal compartments and four cellular-density subregions: tumor with high cellular density (tumor HD), tumor with low cellular density (tumor LD), stroma with high cellular density (stroma HD), and stroma with low cellular density (stroma LD).

The results indicate the presence of obesity-associated morphologic phenotypes in both tumor and stromal compartments, with more pronounced associations observed in stromal regions (stroma HD and stroma LD) than in tumor regions. No distinct subgroup-specific patterns were identified among postmenopausal or ER+/HER2− patients, suggesting that obesity-associated morphologic signatures may be broadly distributed across the examined subgroups.

To further characterize regional variability, we quantified inter-patient heterogeneity using vector embeddings derived from each region of interest. For each region, pairwise cosine distances between slide-level embedding vectors were computed to capture differences in learned morphologic representation. Per-patient average distances were summarized and modeled within each BMI group using kernel density estimation to obtain probability density functions representing inter-patient tumor and stromal heterogeneity. Across all regions, higher BMI was associated with lower average cosine distances, indicating reduced inter-patient morphologic heterogeneity in both tumor and stromal compartments.

## 2 Results

### 2.1 Patient Characteristics

The clinicopathological characteristics of the 195 patients are summarized in Table 1. Of these, 102 (52%) and 93 (48%) were classified into the high- and low-BMI groups, respectively. Overall, 144 patients (74%) were older than 50 years, 166 (85%) were ER-positive, 141 (72%) were PR-positive, 167 (86%) were HER2-negative, and 153 (78%) were diagnosed with invasive ductal carcinoma (IDC). The P-values reported in Table 1 were derived from tests of homogeneity of proportions. No statistically significant differences in the proportions of any reported variable were observed between the high- and low-BMI groups. The magnitude of the differences was also relatively small, with absolute differences of 12 percentage points or less for all variables except age. The proportion of patients older than 50 years differed from 83% in the low-BMI group to 63% in the high-BMI group.

**Table 1:** Clinical characteristics of high and low BMI patients used in this study. P-Values: χ^2^ test for homogeneity (uncorrected), except Age (two-sample Student’s t-test). Statistical tests exclude “Missing” category.

| Clinical characteristic | Value | Overall | High BMI | Low BMI | P-Value |
|---|---|---|---|---|---|
| n |  | 195 | 102 | 93 |  |
| Age, mean (SD) |  | 59.3 (13.4) | 61.7 (12.6) | 56.7 (13.8) | 0.008 |
| Age Class, n (%) | ≥ 50 | 144 (73.8) | 85 (83.3) | 59 (63.4) | 0.003 |
|  | < 50 | 51 (26.2) | 17 (16.7) | 34 (36.9) |  |
| ER, n (%) | Negative | 29 (14.9) | 19 (18.6) | 10 (10.8) | 0.18 |
|  | Positive | 166 (85.1) | 83 (81.4) | 83 (89.2) |  |
| Grade, n (%) | 1 | 56 (28.7) | 27 (26.5) | 29 (31.2) | 0.169 |
|  | 2 | 76 (39.0) | 36 (35.3) | 40 (43.0) |  |
|  | 3 | 61 (31.3) | 38 (37.2) | 23 (24.7) |  |
|  | missing | 2 (1.0) | 1 (1.0) | 1 (1.1) |  |
| HER2, n (%) | Negative | 167 (85.6) | 83 (81.4) | 84 (90.3) | 0.115 |
|  | Positive | 28 (14.4) | 19 (18.6) | 9 (9.7) |  |
| Histology, n (%) | IDC | 153 (78.4) | 82 (80.4) | 71 (76.3) | 0.214 |
|  | ILC | 37 (19.0) | 15 (14.7) | 22 (23.7) |  |
|  | missing | 5 (2.6) | 5 (4.9) | 0 (0.0) |  |
| M, n (%) | 0 | 189 (96.9) | 99 (97.0) | 90 (96.8) | 1 |
|  | 1 | 2 (1.0) | 1 (1.0) | 1 (1.1) |  |
|  | missing | 4 (2.1) | 2 (2.0) | 2 (2.1) |  |
| N, n (%) | 0 | 96 (49.2) | 56 (54.9) | 40 (43.0) | 0.397 |
|  | 1 | 64 (32.8) | 30 (29.4) | 34 (36.6) |  |
|  | 2 | 10 (5.1) | 4 (3.9) | 6 (6.45) |  |
|  | 3 | 11 (5.7) | 5 (4.9) | 6 (6.45) |  |
|  | missing | 14 (7.2) | 7 (6.9) | 7 (7.5) |  |
| PR, n (%) | Negative | 54 (27.7) | 32 (31.4) | 22 (23.7) | 0.297 |
|  | Positive | 141 (72.3) | 70 (68.6) | 71 (76.3) |  |
| T, n (%) | 1 | 108 (55.4) | 53 (52.0) | 55 (59.1) | 0.267 |
|  | 2 | 65 (33.3) | 40 (39.2) | 25 (26.9) |  |
|  | 3 | 17 (8.7) | 7 (6.8) | 10 (10.75) |  |
|  | 4 | 3 (1.6) | 1 (1.0) | 2 (2.15) |  |
|  | missing | 2 (1.0) | 1 (1.0) | 1 (1.1) |  |

### 2.2 Obesity-Associated Morphologic Signatures Are Present in Tumor and Stroma, with a Stronger Signal in Stroma

Morphology-based vector representations were used to build machine learning models that classify high-vs low-BMI patients from tumor, stroma, combined tumor-stroma, and subregions based on cellular density. Figure 2 summarizes performance results for different model types using 5×2 cross-validation, with corresponding numerical results provided in Supplementary Table S1.

Most classifiers achieved median AUC values greater than 0.5, and with a few exceptions in specific tissue regions, classifiers were statistically significant at the 5% level. In general, logistic regression classifiers generated higher performance scores compared to other models. Among the evaluated tissue categories, stromal regions yielded the highest classification performance, with a mean AUC of 0.78 (95% CI: 0.73–0.83) when using logistic regression. Similar performance was observed for stroma LD (AUC = 0.78, 95% CI: 0.74–0.82) and stroma HD (AUC = 0.77, 95% CI: 0.71–0.82) also using logistic regression. In contrast, tumor and combined tumor– stroma regions produced lower AUC values (Figure 2; Supplementary Table S1).

### 2.3 No Evidence of Cohort-Specific Differences in Obesity Signature Detection

Morphology-based vector representations were used to build logistic regression models that classify high-vs low-BMI patients from tumor, stroma, and combined tumor-stroma regions across the different patient cohorts listed in Table 2. Figure 3 summarizes model performance results with corresponding numerical results provided in Supplementary Table S2.

**Table 2:** Selected patient cohorts for analysis.

| **Cohort designation** | **Description** | **Patient Count** |  |  |
|---|---|---|---|---|
|  |  | **Total** | **High BMI** | **Low BMI** |
| ALL PAT | All patients | 195 | 102 | 93 |
| ER+/HER2- | ER+/HER2- patients | 147 | 72 | 75 |
| ALL PAT (50) | Patients with Age ≥ 50 | 144 | 85 | 59 |
| ER+/HER2- (50) | ER+/HER2- patients with Age ≥ 50 | 108 | 62 | 46 |
| DELTA | Patients that are not ER+/HER2- | 48 | 30 | 18 |

Across all cohorts, stromal embeddings consistently yielded higher AUC values than tumor or combined tumor–stroma embeddings. In the ER+/HER2− cohort, the stroma-based model achieved an AUC of 0.74 (95% CI: 0.68–0.79), compared with 0.65 (95% CI: 0.56–0.73) for the tumor-based model. Similarly, in the ALL PAT (50) cohort, the stroma-based model achieved an AUC of 0.70 (95% CI: 0.62–0.79), whereas the tumor-based model achieved an AUC of 0.61 (95% CI: 0.52–0.71). Similar region-based performance trends were observed across the remaining clinically relevant cohorts. Statistical comparison using a two-sample Student’s t-test of stroma- and tumor-based AUCs (Table 3) showed significant differences for the ALL PAT and ER+/HER2− cohorts (p < 0.05).

**Table 3:**
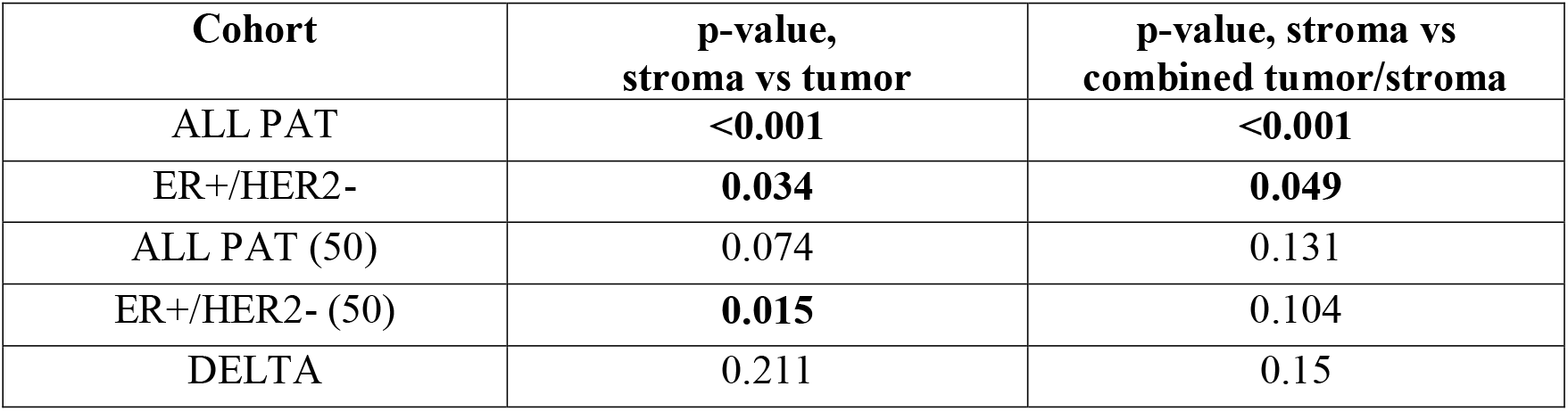
P-values from two-sample t-tests comparing mean logistic regression AUC values for BMI classification across stromal, tumor, and combined tumor–stroma regions, evaluated separately within each cohort.

### 2.4 Higher BMI Is Associated with Reduced Inter-Patient Morphologic Heterogeneity in Tumor and Stromal Regions

Morphology-based representation vectors from tumor, stroma and subregions based on cellular density were used to quantify inter-patient heterogeneity based on each patient’s average morphologic distance to the remaining patients in the cohort. These average distances were used to generate probability density functions, which were compared across groups to assess differences in inter-patient heterogeneity.

Higher BMI was associated with lower inter-patient heterogeneity in both tumor and stroma compartments (Figure 4, top row); however, a statistically significant difference between BMI groups was observed only for the stromal compartment (Kolmogorov–Smirnov test, Table 4). Similar shifts toward lower heterogeneity were observed in the tumor LD, tumor HD, stroma LD, and stroma HD subregions (Figure 4, middle and bottom rows). Among these, tumor LD, tumor HD, and stroma HD showed significant differences between BMI groups, whereas stroma LD did not (Table 4).

**Table 4:**
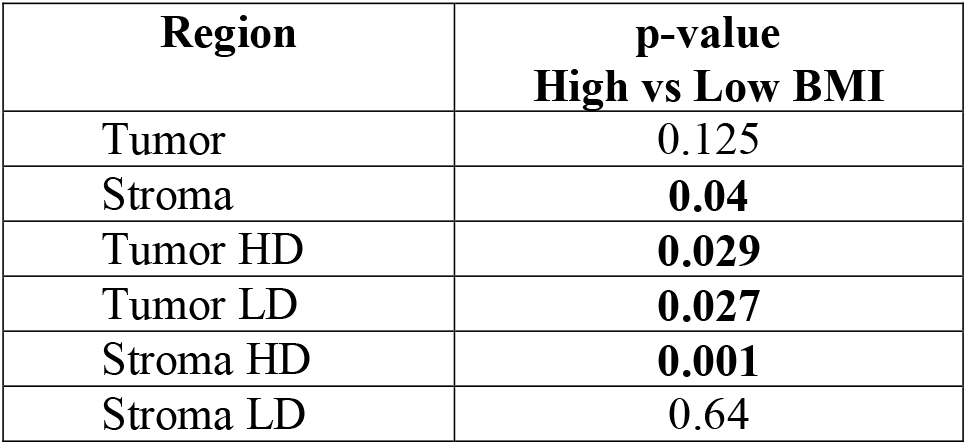
P-values (uncorrected) from Kolmogorov-Smirnov tests comparing inter-patient heterogeneity probability density functions for high and low BMI patients evaluated for the different regions of interest.

## 3 Discussion

The present study investigated whether obesity is associated with detectable morphological alterations in tumor cells and the surrounding stromal compartment captured from routine histopathology images. To address this question, we employed an artificial intelligence foundation model to generate tile- and slide-level representation vectors from digitized tissue sections (Zalach, et al., 2024). Regions of interest were defined as tumor, stroma, and their respective low- and high-cell-density subregions (tumor LD, tumor HD, stroma LD, and stroma HD). A classifier was then trained to distinguish patients with high BMI (≥30) from those with low BMI (<25). Our results demonstrate that obesity signatures are detectable from both stromal and tumor regions, with stromal regions consistently yielding superior classification performance. Notably, we observed a progressive improvement in detectability when moving from tumor HD to tumor LD and, ultimately, to stromal components, suggesting that the morphological imprint of obesity is more pronounced or more readily captured by the model in the stromal compartment.

An important consideration is that the stromal regions analyzed in this study did not include peritumoral adipose tissue. During preprocessing, image tiles containing large adipose areas were excluded using a dedicated neural network trained to filter out tiles in which adipose tissue occupied more than 1/8 of the tile area. While small residual adipose fragments may remain within some tiles and could contribute to the classification signal, the ability to detect obesity-associated signatures within tumor regions, including tumor HD subregions, suggests that residual adipose tissue is unlikely to be the sole source of the observed signal. Notably, predictive performance in tumor HD subregions was generally lower than in stromal compartments, indicating that signal extraction in these regions is more challenging. However, several algorithms, including logistic regression, achieved statistically significant classification results, supporting the presence of detectable obesity-associated features even in tumor-dominant areas.

Together, these findings are consistent with the possibility that obesity-related morphological alterations extend beyond direct adipose representation and may involve broader changes in the tumor and stromal microenvironment. This interpretation aligns with reports indicating that obesity is associated with extracellular matrix remodeling, fibrosis, and immune cell infiltration in the breast stroma (Hawes, et al., 2025; Hillers-Ziemer, et al., 2022; Seo, et al., 2015).

The precise spatial localization of obesity-related signals within these regions remains an open question. Future work could leverage interpretability methods such as those that construct heatmaps of attention coefficients (Ilse, et al., 2018; Albuquerque, et al., 2024), or those that synthesize image counterfactuals to provide explainability (Kaczmarzyk, et al., 2024). These methods identify the specific tissue structures and cellular neighborhoods that contribute most strongly to the classification. Such analyses could help determine whether the signal arises from specific stromal cell populations, extracellular matrix features, immune cell infiltrates, or combinations of these and other elements.

We also evaluated obesity signature detection across clinically relevant patient subgroups, including subgroups defined by estrogen-receptor status and age. Similar to the full cohort, stromal-based models consistently outperformed tumor-based models in all subgroups, with no substantial differences observed between subgroups. However, the smaller sample sizes increased performance variability and may have limited the power to detect subgroup-specific effects. Given the established heterogeneity in obesity–breast cancer associations across molecular subtypes and menopausal status (Reeves, et al., 2007; Munsell, et al., 2014; García-Estévez, et al., 2021; Lohmann, et al., 2021; Sparano, et al., 2012) larger, stratified cohorts will be needed to determine whether obesity-associated histologic signatures differ among these clinically relevant populations.

Beyond the detection of obesity signatures, the AI-derived representation vectors offer a powerful framework for quantifying inter-patient morphological heterogeneity. By computing pairwise dissimilarities among patients’ region-based slide-level embeddings, we developed distributions of inter-patient heterogeneity for each region of interest. This approach enables a data-driven comparison of morphological diversity between BMI groups that does not rely on predefined histopathological or molecular features.

Our analysis revealed a statistically significant reduction in inter-patient stromal heterogeneity among high-BMI patients compared to low-BMI patients. This finding suggests that obesity may impose a degree of morphological convergence on the stromal microenvironment potentially reflecting obesity-driven remodeling processes such as fibrosis, immune infiltration patterns, or extracellular matrix reorganization (Seo, et al., 2015; Hillers-Ziemer, et al., 2022; Hawes, et al., 2025; Yang, et al., 2023). In contrast, the difference in inter-patient tumor heterogeneity between BMI groups, although visually apparent in the probability density estimates, did not reach statistical significance when the tumor region was analyzed as a whole.

When the tumor compartment was subdivided into tumor LD and tumor HD subregions, the differences in inter-patient heterogeneity between BMI groups reached statistical significance. This observation may be related to the region selection procedure. As described in the Methods section, the clustering approach used to define LD and HD categories excludes tiles with ambiguous assignments, thereby restricting the analysis to tiles with more distinct morphological characteristics. Such filtering alters the tile composition within each subgroup and could contribute to the detection of between-group differences. However, this process may also reduce the ability of a model to differentiate high and low BMI, which could contribute to the non-significant difference observed for stroma LD.

These findings have methodological implications for future studies of tumor heterogeneity using AI-based embeddings. The observation that curated subregions yield stronger statistical signals than broadly defined compartments suggests that deliberate region refinement, whether based on cell density, as performed here, or on other tissue characteristics can enhance the sensitivity of heterogeneity analyses. For example, isolating regions enriched for specific cell populations such as cancer-associated fibroblasts (Yang, et al., 2023; Sahai, et al., 2020) or tumor-infiltrating lymphocytes (Ciarka, et al., 2024; Finkelman, et al., 2025) and repeating the inter-patient heterogeneity analysis could reveal additional associations with BMI or other clinical variables. Moreover, these curated subregions could be used in conjunction with obesity signature detection to determine whether specific cellular neighborhoods harbor identifiable morphological changes linked to obesity.

Taken together, the two lines of investigation presented in this study, obesity signature detection and inter-patient heterogeneity analysis, converge on the conclusion that obesity leaves a measurable morphological imprint on both tumor and stromal tissue. The stromal compartment appears to be particularly informative, both as a substrate for obesity classification and as a site of reduced inter-patient diversity in the setting of high BMI. These results underscore the potential of AI-driven histologic analyses to complement traditional epidemiological and molecular approaches in elucidating the complex relationship between obesity and breast cancer biology.

Finally, we summarize the main limitations of the current study, some of which have been discussed throughout the manuscript. The relatively small, single-institution cohort of 195 breast cancer patients limits the generalizability of the models and highlights the need for validation in larger, multi-institutional cohorts. Clinically defined subgroups, including ER+/HER2− and age-stratified groups, had smaller sample sizes and reduced statistical power and should therefore be considered exploratory. Cell-density regions were defined using unsupervised clustering of tile vectors, providing only an approximate characterization of tissue cellularity. Future studies incorporating cell segmentation and direct cell counts may provide a more accurate characterization of cellular density.

## 4 Methods

### 4.1 Ethics statement

This retrospective study conducted at the Ellison Medical Institute and Walter Reed National Military Medical Center was approved by Western Institutional Review Board-Copernicus Group (IRB Exemption) and Walter Reed National Military Medical Center Institutional Review Board (WRNMMC-2019-0228). All research was performed in accordance with relevant guidelines and regulations.

### 4.2 Study participants and patient cohorts

A total of 337 H&E-stained breast cancer tissue sections from approximately the same number of patients were obtained from the Murtha Cancer Center at Walter Reed National Military Medical Center, together with associated clinical data, including body mass index (BMI) and other relevant variables. After quality control, duplicate slides from patients with multiple available sections were removed. Slides from male patients and patients with incomplete clinical data were also excluded, resulting in a curated dataset of 304 whole-slide images (WSIs).

The slides were then stratified by patient BMI into high-BMI (BMI ≥ 30; n = 102) and low-BMI (BMI < 25; n = 93). Patients with intermediate BMI values (25 ≤ BMI < 30) were excluded to maximize the contrast between groups and facilitate training of the machine learning (ML) classifiers.

The primary analysis was conducted using the complete cohort of 195 breast cancer patients (ALL PAT), with secondary analyses performed on clinically relevant subgroups. Table 2 summarizes the analyzed cohorts and their corresponding sample sizes. The ER+/HER2− cohort includes patients with estrogen receptor-positive and HER2-negative breast cancer. The ALL PAT (50) cohort includes all patients aged 50 years or older and was used as an approximate representation of a postmenopausal population. The ER+/HER2− (50) cohort further restricts the ER+/HER2− group to patients aged 50 years or older. Finally, the DELTA cohort includes patients who were not classified as ER+/HER2−. A cohort representing premenopausal patients was not included because of the insufficient number of eligible cases.

### 4.3 Acquisition of H&E-Stained Histopathology Images

Slides were digitized using a VS200 Olympus scanner at 20x magnification (0.274 μm/pixel) and saved as uncompressed images to preserve maximum quality.

Multiple rounds of visual quality control were performed to minimize scanning artifacts and identify slides with potential sources of error for downstream image processing and machine learning workflows. WSIs exhibiting significant degradation—including tissue folds, foreign artifacts (e.g., dust, bubbles), coverslip marks, excessive tissue fragmentation, or poor staining quality—were excluded from the dataset.

### 4.4 Image Preprocessing

WSIs were processed according to the workflow outlined in Figure 1A. Each slide was first divided into a grid of 380 × 380–pixel tiles, a dimension optimized for input into the CanvOI foundation model.

**Figure 1.**
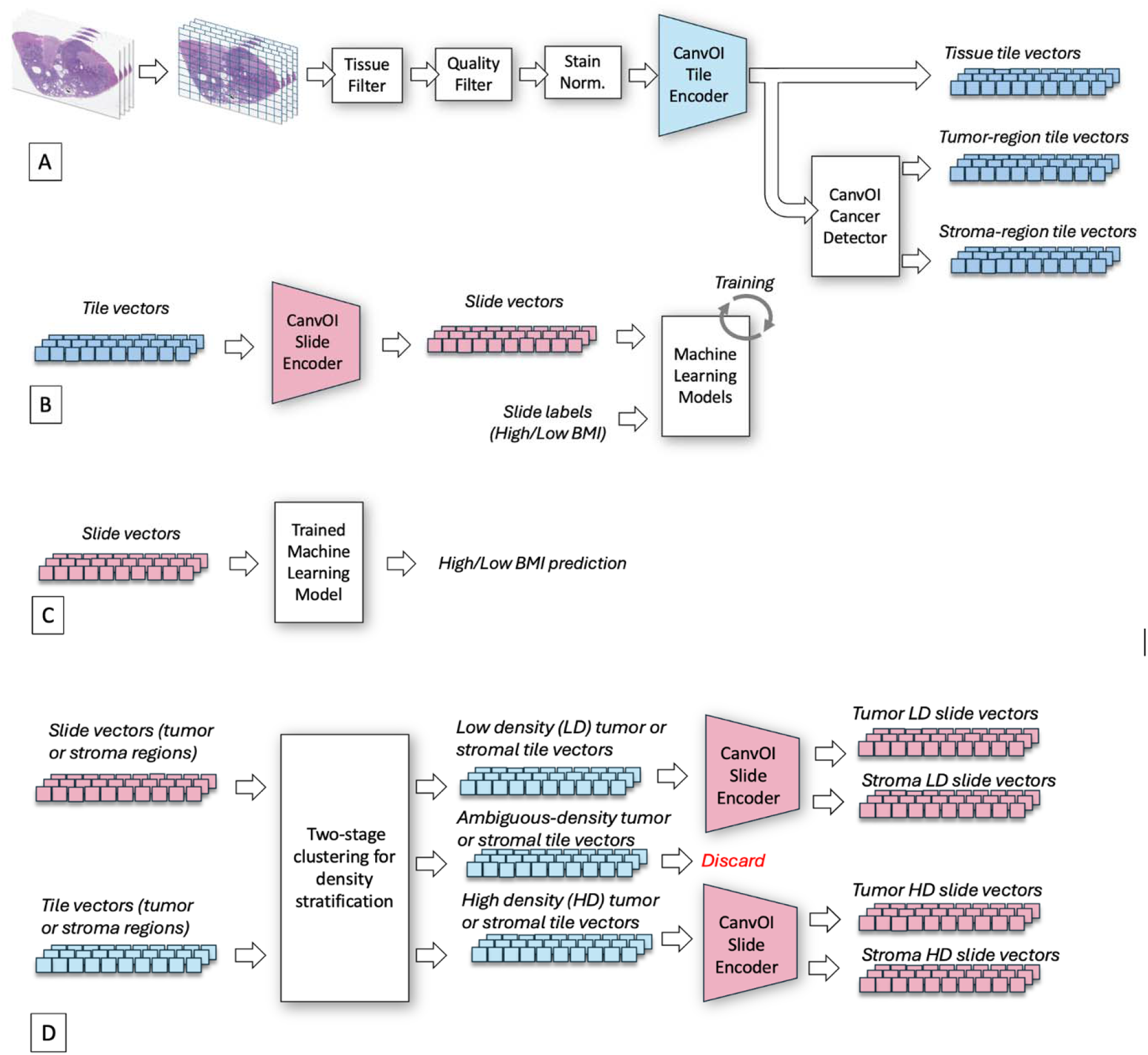
Data processing workflows in this study. A: Pre-processing of whole slide images to generate tiles and tile-level vector embeddings for the tumor, stroma, and combined tumor/stroma regions. B: Aggregation of tile vectors to generate slide-level vector embeddings. These vectors and the matching slide labels are used for training task-specific machine-learning models. C: Slide vectors from test whole-slide images become inputs to the trained models to obtain class predictions for inference and performance evaluation. D: Slide- and tile-level vectors extracted from tumor and stromal regions are used in a clustering algorithm to classify tiles according to low or high cellular density. Tile-level vectors within each density-defined subregion are then aggregated to generate slide-level vectors, which, together with slide-level labels, are used for machine learning–based BMI prediction and inter-patient heterogeneity analysis.

**Figure 2.**
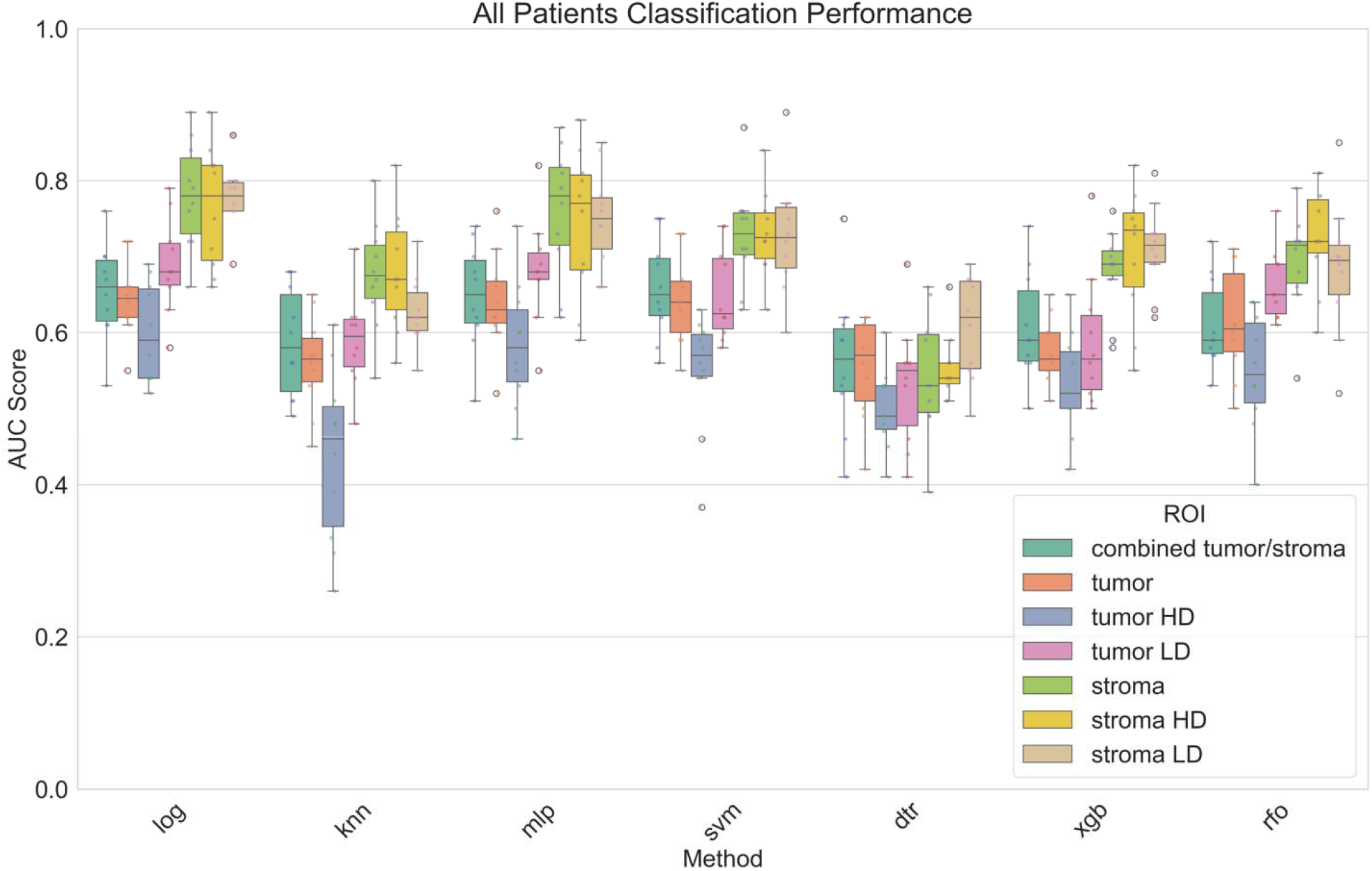
Area Under the Curve (AUC) values obtained from 5×2 cross-validation for multiple machine learning classifiers trained to distinguish high- and low-BMI patients using vector representations derived from combined tumor–stroma, tumor, stroma, tumor LD, tumor HD, stroma LD, and stroma HD regions. Results are shown for the complete patient cohort. Each boxplot summarizes the distribution of AUCs across ten experiments. The central line indicates the median; box edges represent the third (Q3) and first (Q1) quartiles. Whiskers extend to the most extreme values within 1.5 × the interquartile range (IQR), and open circles denote outliers beyond this range.

**Figure 3.**
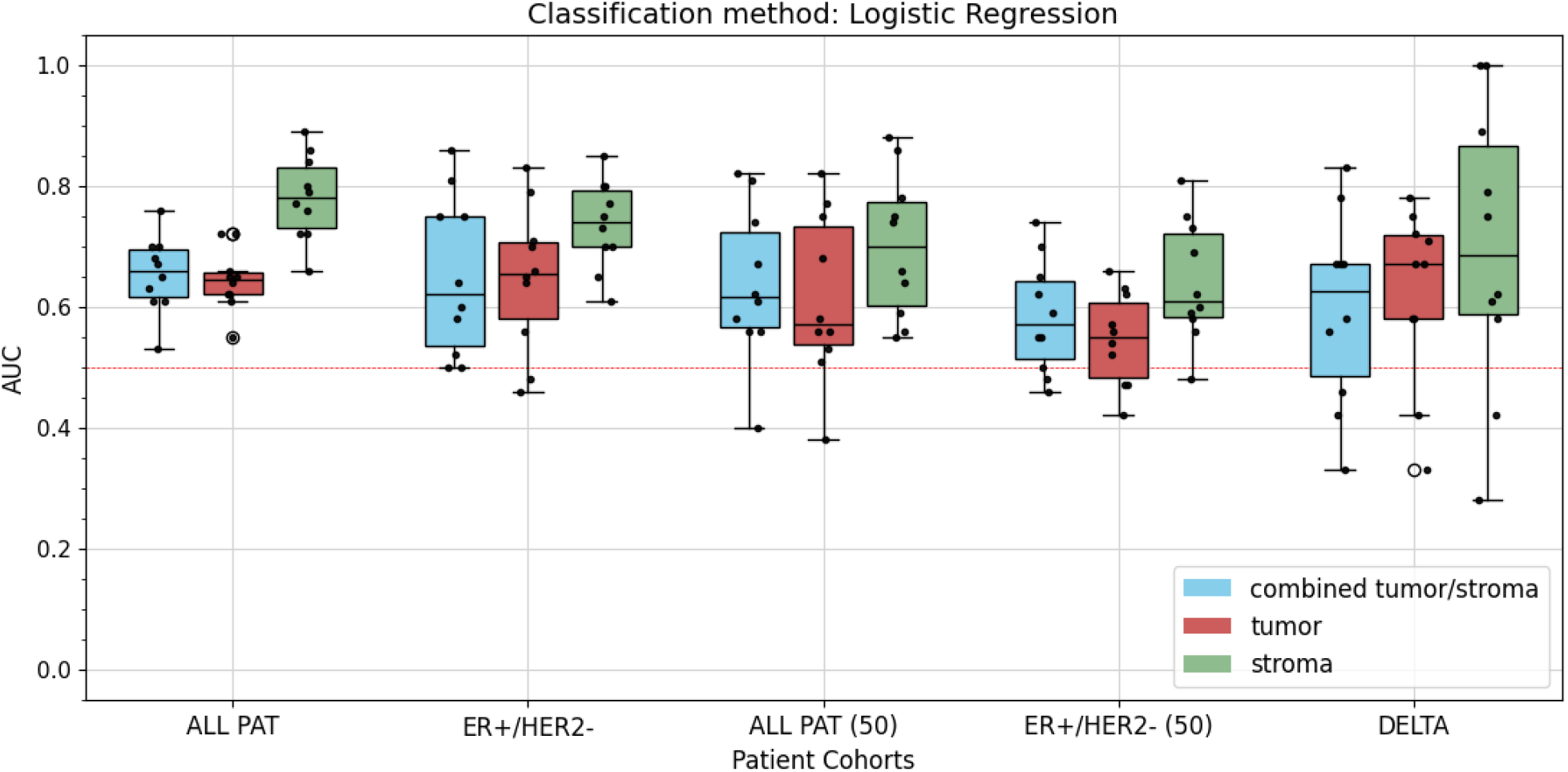
Area under the curve (AUC) values obtained from 5×2 cross-validation using logistic regression to distinguish high- and low-BMI patients using vector representations derived from tumor, stromal, and combined tumor–stroma regions. Results are shown for the following cohorts: all patients, ER+/HER2− patients, patients older than 50 years, ER+/HER2− patients older than 50 years, and patients not classified as ER+/HER2− (DELTA). Each boxplot summarizes the distribution of AUCs across ten experiments. The central line indicates the median; box edges represent the third (Q3) and first (Q1) quartiles. Whiskers extend to the most extreme values within 1.5 × the interquartile range (IQR), and open circles denote outliers beyond this range.

**Figure 4.**
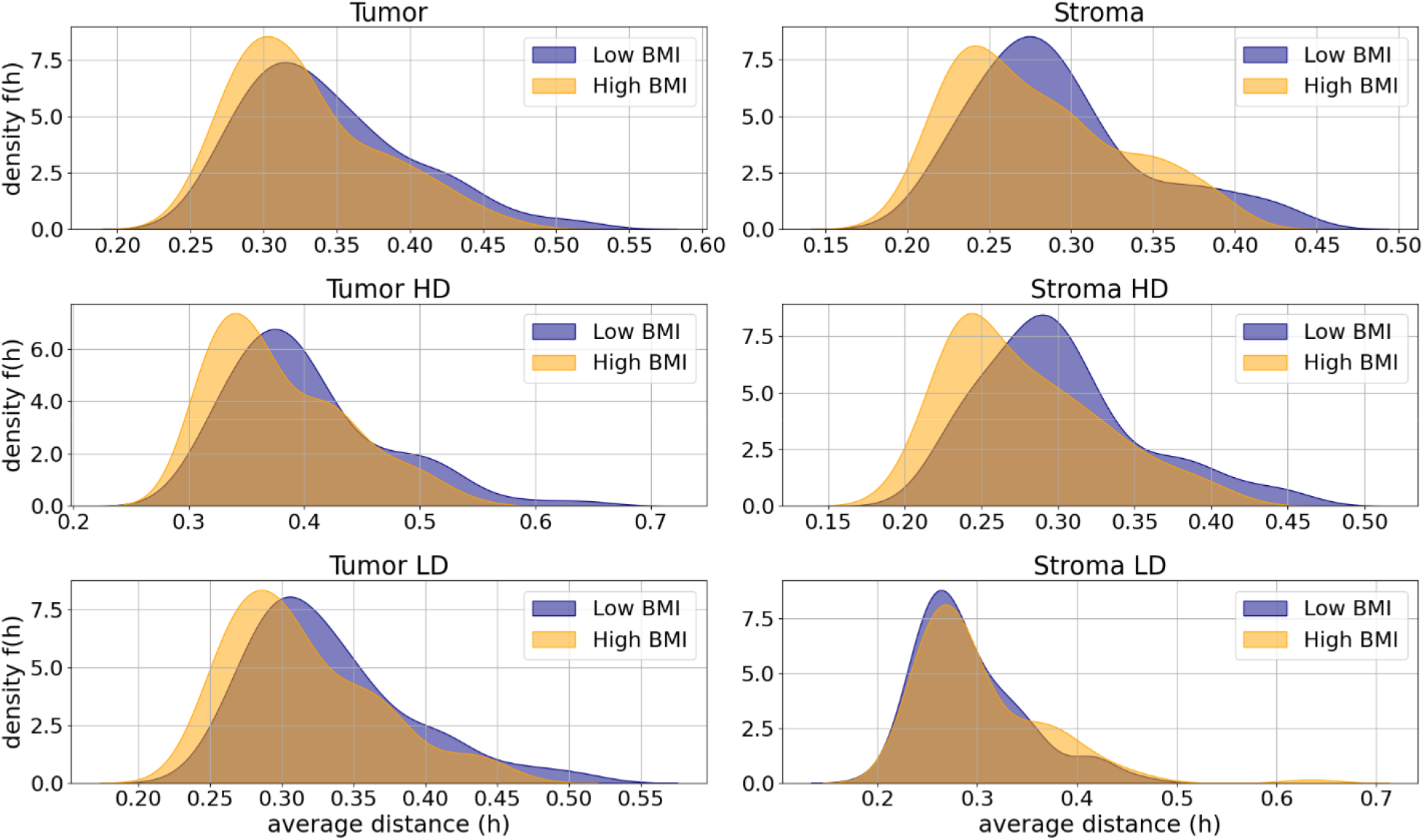
Inter-patient heterogeneity density functions for different regions stratified by BMI category (high vs. low). Top row: density functions extracted from tumor (left) and stromal (right) regions; middle and bottom rows: density functions derived from tumor and stroma subregions with high cellular density (HD) and low cellular density (LD), respectively.

To exclude tiles containing little or no relevant tissue, a ResNet-50 convolutional neural network (Tissue Filter; Figure 1A) was trained to distinguish tissue-containing from non-tissue tiles. The positive training subset comprised thousands of tiles containing at least 7/8 of visible tumor or stromal tissue, whereas the negative subset comprised thousands of tiles containing at least 7/8 visible adipose or non-tissue areas. Training labels were assigned by visual assessment to ensure compliance with these criteria. Representative examples of retained and discarded tiles are shown in Supplementary Figure S1.

Following tissue filtering, two additional ResNet-50 networks were applied for quality control (collectively referred to as the Quality Filter in Figure 1A). The first model identified and removed tiles with significant out-of-focus artifacts, and the second excluded tiles with excessive ink markings. The remaining tiles underwent stain normalization using a modified Reinhard algorithm (Roy, et al., 2021), selected on the basis of qualitative visual assessment showing superior performance relative to other commonly used normalization techniques.

### 4.5 Generating Representation Vectors Using the CanvOI Foundation Model

The CanvOI foundation model, operating as a tile encoder, generates a 1536-dimensional representation vector for each image tile (Figure 1A). These tile-level vectors also serve as input to a CanvOI-based pan-cancer detector, which classifies each tile as tumor or non-tumor. Supplementary Figure S2 presents a representative breast cancer WSI from the BACH dataset with tumor regions annotated by expert pathologists (Aresta et al., 2019), alongside an overlay of tumor tiles identified by the pan-cancer detector.

The workflow generates three primary tile-vector sets corresponding to combined tumor–stroma, tumor-only, and stromal regions, which are analyzed through independent downstream pipelines (Figures 1B and 1D). In slide-encoding mode (Figure 1B), the CanvOI foundation model aggregates tile-level vectors into 2000-dimensional slide-level embeddings for each tissue compartment. These embeddings are used as inputs for BMI classification models and inter-patient heterogeneity analysis. For cellularity-based analysis (Figure 1D), tumor and stromal tiles are further partitioned by clustering into four subregions: tumor with low cellular density (tumor LD), tumor with high cellular density (tumor HD), stroma with low cellular density (stroma LD), and stroma with high cellular density (stroma HD). Slide-level embeddings from these subregions are similarly evaluated for BMI prediction and heterogeneity analysis.

### 4.6 Machine Learning Classifiers: Training and Evaluation

Slide-level embeddings and corresponding BMI labels were used to train machine learning classifiers to identify associations between encoded tissue morphology and BMI status. The evaluated models included logistic regression (log), k-nearest neighbors (knn), multilayer perceptron (mlp), support vector machines (svm), decision trees (dtr), XGBoost (xgb), and random forest (rfo) classifiers. All models were implemented using default hyperparameters from the Python scikit-learn and XGBoost libraries to enable standardized comparisons.

Model performance was evaluated using held-out WSIs through a 5×2 cross-validation scheme consisting of five-fold cross-validation repeated twice with different random seeds. In each replicate, 80% of samples were used for training and 20% for testing, with identical data partitions maintained across experiments. Predictions were compared with ground-truth BMI labels using the area under the receiver operating characteristic curve (AUC).

An exploratory analysis incorporating overweight patients (25 ≤ BMI < 30) as an additional test group did not achieve acceptable classification performance, suggesting that the models may preferentially capture stronger obesity-associated signals rather than subtler morphological patterns associated with intermediate BMI.

### 4.7 Unsupervised Clustering for Tile-Level Cellular Density Stratification

Tumor and stromal regions exhibit variation in cell density that reflects distinct biological features. In tumors, increased cellular density may indicate proliferative activity and aggressive disease, while in stroma, cellular density reflects composition such as lymphocyte infiltration or abundance of cancer-associated fibroblasts.

To characterize these differences, we applied a machine learning–based clustering approach to partition tumor and stromal tiles according to cellular density. The method groups tile embeddings into visually distinguishable cellularity-based clusters, providing an approximate tissue stratification rather than a direct measurement of cell density. Future studies incorporating cell segmentation or direct cell-count measurements may enable more precise characterization.

Separating image tiles into low- and high-cellular density groups for tumor and stromal regions required a purpose-built clustering algorithm applied to tile-level vector embeddings extracted with a CanvOI encoder. Embedding dimensionality was reduced from 1536 to 50 using principal component analysis (PCA). The algorithm follows a two-stage procedure: initialization and expansion.

During initialization, a small set of slide-level tumor images (n=4) was selected, including two with the largest and two with the smallest slide-level vector differences. Tile-level vectors from these tumor images were input into a Gaussian Mixture Model (GMM) to generate two clusters and their corresponding centroids. Visual inspection indicated that the centroids separated tiles primarily by cell density.

In the expansion phase, the full collection of tumor tiles was clustered using the initialized centroids and the dimensionality-reduced embeddings. Updated centroids were obtained, and each tile was assigned a probability of membership. Tiles with ≥70% probability for either centroid were retained, while the remainder were discarded. The resulting clusters approximately correspond to tumor with low cell density (tumor LD) and tumor with high cell density (tumor HD).

The same two-stage procedure was applied to stromal regions. For initialization, seven stromal images were selected based on extreme slide-level vector embeddings (three with maximum slide-level vector differences and four with minimum differences). Their tiles were used to generate initial centroids, after which all stromal tiles were clustered. This process produced stromal tiles approximately classified as stroma with low cell density (stroma LD) and with high cell density (stroma HD).

Figure 1D shows a diagram describing the clustering process using tile and slide vectors. Supplementary figures S3 and S4 show examples of tiles grouped into HD and LD groups according to the algorithm described in this section.

### 4.8 Inter-tumor and inter-stroma heterogeneity density measurements using vector embeddings

Large-scale foundation models like CanvOI produce vector embedding spaces that organize according to tissue morphology and disease state, as evidenced by clustering behavior and strong linear probing performance (Caron, et al., 2021; Zalach, et al., 2024). Accordingly, distances between embedding vectors can be used to quantify homogeneity and heterogeneity across patient tissue samples.

Let *R* denote a region of interest within a tissue sample. In this study, *R* corresponds to tumor, stroma, tumor LD, tumor HD, stroma LD, or stroma HD. Each region *R* is composed of a collection of tiles, each associated with a tile-level vector embedding. These tile-level embeddings are aggregated into a slide-level vector *x*^*R*^ using a slide-encoder module (e.g., CanvOI’s slide encoder).

Given two patients with regional vectors 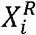 and 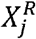, their semantic dissimilarity 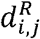 is computed using the cosine distance:

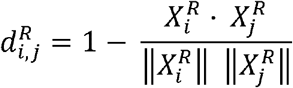

where the numerator denotes the dot product and the denominator the product of vector norms. Cosine distance ranges from 0 (identical orientation) to 2 (maximally opposite orientation).

For a cohort of *K* patients, pairwise dissimilarities are assembled into a symmetric matrix *D*^*R*^ ∈ ℝ^*K* × *K*^ where each entry corresponds to 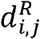. Computing the per-row (or equivalently per-column) mean yields a *K*-dimensional vector *H*^*R*^, whose elements (*h*_1_,*h*_2_,…, *h*_K_) represent the average distance of each patient relative to all others in the cohort. The per-patient average distance *h*_*q*_ constitutes a measure of inter-patient heterogeneity:

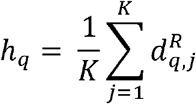

Treating *H*^*R*^ as a set of observations, we apply kernel density estimation to obtain a probability density function *f*^*R*^ (*h*), which characterizes the distribution of inter-patient heterogeneity values within the cohort. A density function concentrated at lower *h* values represents greater homogeneity, whereas concentration at higher values reflects increased heterogeneity.

For each region of interest *R*, we compare the probability density curves *f*^*R*^ (*h*) between the high- and low-BMI groups. Statistical differences between density functions are assessed using the Kolmogorov–Smirnov (KS) test.

## 5 Data availability

The original whole-slide images (WSIs) are not publicly available due to patient privacy and data protection policies. To facilitate reproducibility and support further research, the WSIs were processed to identify tissue regions, which were subsequently divided into non-overlapping tiles of 380 × 380 pixels. Vector embeddings were extracted from these tiles using the CanvOI Pathology Foundation Model. In addition to the tile-level embeddings, vector representations of tumor and stroma regions, generated using the CanvOI slide encoder, are also provided. The dataset, comprising the vector embeddings and associated body mass index (BMI) information, is publicly available through Zenodo at https://doi.org/10.5281/zenodo.23992255.

## 6 Code availability

The pre-processing software used for quality control, tissue filtering, and stain normalization is proprietary software owned by the Ellison Medical Institute. The CanvOI tile and slide encoders, as well as the cancer detector, are proprietary technologies owned by Imagene AI. The code used for vector analysis was developed in Python and comprises scripts and Jupyter notebooks implementing machine learning, statistical analysis, clustering and data visualization methods. The complete vector analysis codebase is publicly available in the following GitHub repository: https://github.com/eitm-org/breast-cancer-bmi-signatures.

## 7 Acknowledgements

This research was partially supported by a generous grant from the Breast Cancer Research Foundation (BCRF-21-002). Additional funding for these efforts was provided from Uniformed Services University of Health Sciences (USUHS) award from the Defense Health Program to the Murtha Cancer Center Research Program (HU00012320028, JSH Lee/DB Agus) administered by the Henry M. Jackson Foundation for the Advancement of Military Medicine. Imagene AI provided access to the CanvOI digital pathology foundation model.

## 8 Disclaimer

The contents of this document are the sole responsibility of the authors and do not necessarily reflect the views, assertions, opinions, or policies of the Uniformed Services University of the Health Sciences, the Department of War, or the Departments of the Army, Navy, or Air Force. Mention of trade names, commercial products, or organizations does not imply endorsement by the U.S. government.

The view(s) expressed herein are those of the author(s) and do not reflect the official policy or position of the Henry M. Jackson Foundation for the Advancement of Military Medicine, Inc.

## 9 Author Contributions

D.B.A, N.M., R.B., J.S.H.L., J.L., J.W. conceptualized the study and supervised research and project resources. J.L., I.T.L., T.J.J., O.C., B.T. collaborated on data collection, data exchange, and policy compliance. B.S., B.T., D.S. contributed to specimen management and scanning. S.P., L.P., J.P., T.K., Z.F., B.S. contributed to performing slide quality control. J.L. and I.T.L. contributed to pathology interpretations. R.B. and V.H.N. contributed to clinical data interpretations and data standardizations. N.M., T.J.J., R.B. E.A.H., S.P. supervised overall project development and research methodology development. E.A.H., S.P., L.P, J.P., T.K., Z.F. I.K. developed the AI/ML software pipeline, research methods, and contributed to formal data analysis. All authors contributed to drafting the manuscript and provided critical revisions. Large language models (LLMs) were used solely to refine the language and clarity of the manuscript. They were not used to generate scientific outputs, influence scientific interpretations, or make scientific judgements. All scientific content, interpretations, and conclusions were developed and validated by the authors.

