## Supplemental figures and tables for "Deep Learning Identifies BMI-Associated Morphologic Features in Breast Cancer Tissue"

### Supplementary Material

Table S1: Performance metrics for high versus low BMI classification across different machine learning methods and tissue regions. Models were trained and evaluated using 5×2 cross-validation on the full patient cohort. Reported p-values, obtained from a one-sample t-test, assess whether the mean AUC exceeds the performance of a random classifier (AUC = 0.5).

| **Method** | **Region** | **Mean AUC** | **Median AUC** | **95% CI** | **P value** |
| --- | --- | --- | --- | --- | --- |
| log | combined tumor/stroma | 0.65 | 0.66 | [0.61, 0.7] | <0.001 |
|  | tumor | 0.65 | 0.65 | [0.61, 0.68] | <0.001 |
|  | stroma | 0.78 | 0.78 | [0.73, 0.83] | <0.001 |
|  | tumor HD | 0.6 | 0.59 | [0.55, 0.65] | <0.001 |
|  | tumor LD | 0.69 | 0.68 | [0.64, 0.73] | <0.001 |
|  | stroma HD | 0.77 | 0.78 | [0.71, 0.82] | <0.001 |
|  | stroma LD | 0.78 | 0.78 | [0.74, 0.82] | <0.001 |
| knn | combined tumor/stroma | 0.59 | 0.58 | [0.54, 0.64] | 0.002 |
|  | tumor | 0.56 | 0.56 | [0.51, 0.61] | 0.007 |
|  | stroma | 0.68 | 0.68 | [0.62, 0.73] | <0.001 |
|  | tumor HD | 0.44 | 0.46 | [0.36, 0.52] | 0.94 |
|  | tumor LD | 0.59 | 0.59 | [0.55, 0.63] | <0.001 |
|  | stroma HD | 0.68 | 0.67 | [0.62, 0.74] | <0.001 |
|  | stroma LD | 0.63 | 0.62 | [0.59, 0.66] | <0.001 |
| mlp | combined tumor/stroma | 0.65 | 0.65 | [0.6, 0.7] | <0.001 |
|  | tumor | 0.64 | 0.63 | [0.59, 0.69] | <0.001 |
|  | stroma | 0.76 | 0.78 | [0.7, 0.82] | <0.001 |
|  | tumor HD | 0.58 | 0.58 | [0.53, 0.64] | 0.005 |
|  | tumor LD | 0.68 | 0.68 | [0.63, 0.73] | <0.001 |
|  | stroma HD | 0.74 | 0.77 | [0.67, 0.81] | <0.001 |
|  | stroma LD | 0.75 | 0.75 | [0.7, 0.8] | <0.001 |
| svm | combined tumor/stroma | 0.66 | 0.65 | [0.61, 0.7] | <0.001 |
|  | tumor | 0.64 | 0.64 | [0.6, 0.68] | <0.001 |
|  | stroma | 0.73 | 0.73 | [0.68, 0.78] | <0.001 |
|  | tumor HD | 0.55 | 0.57 | [0.49, 0.61] | 0.04 |
|  | tumor LD | 0.65 | 0.62 | [0.61, 0.69] | <0.001 |
|  | stroma HD | 0.73 | 0.72 | [0.69, 0.77] | <0.001 |
|  | stroma LD | 0.73 | 0.72 | [0.67, 0.78] | <0.001 |
| dtr | combined tumor/stroma | 0.56 | 0.56 | [0.49, 0.63] | 0.033 |
|  | tumor | 0.55 | 0.57 | [0.51, 0.6] | 0.015 |
|  | stroma | 0.54 | 0.53 | [0.49, 0.6] | 0.063 |
|  | tumor HD | 0.50 | 0.49 | [0.46, 0.54] | 0.523 |
|  | tumor LD | 0.53 | 0.55 | [0.48, 0.59] | 0.109 |
|  | stroma HD | 0.55 | 0.54 | [0.52, 0.59] | 0.002 |
|  | stroma LD | 0.61 | 0.62 | [0.56, 0.66] | <0.001 |
| xgb | combined tumor/stroma | 0.61 | 0.59 | [0.56, 0.66] | <0.001 |
|  | tumor | 0.58 | 0.56 | [0.55, 0.61] | <0.001 |
|  | stoma | 0.68 | 0.69 | [0.64, 0.72] | <0.001 |
|  | tumor HD | 0.53 | 0.52 | [0.48, 0.58] | 0.092 |
|  | tumor LD | 0.59 | 0.56 | [0.53, 0.65] | 0.005 |
|  | stroma HD | 0.71 | 0.73 | [0.64, 0.77] | <0.001 |
|  | stroma LD | 0.71 | 0.71 | [0.67, 0.75] | <0.001 |
| rfo | combined tumor/stroma | 0.61 | 0.59 | [0.57, 0.65] | <0.001 |
|  | tumor | 0.61 | 0.60 | [0.56, 0.66] | <0.001 |
|  | stoma | 0.69 | 0.71 | [0.64, 0.74] | <0.001 |
|  | tumor HD | 0.55 | 0.55 | [0.49, 0.6] | 0.038 |
|  | tumor LD | 0.66 | 0.65 | [0.63, 0.7] | <0.001 |
|  | stroma HD | 0.73 | 0.72 | [0.68, 0.78] | <0.001 |
|  | stroma LD | 0.68 | 0.69 | [0.62, 0.75] | <0.001 |

Table S2: Performance metrics for high versus low BMI classification across different cohorts and tissue regions using a logistic regression classifier. Models were trained and evaluated using 5×2 cross-validation. Reported p-values, obtained from a one-sample t-test, assess whether the mean AUC exceeds the performance of a random classifier (AUC = 0.5).

| **Region** | **Cohort** | **Mean AUC** | **CI (95%)** | **Median AUC** | **Max AUC** | **Min AUC** | **P value** |
| --- | --- | --- | --- | --- | --- | --- | --- |
| combined tissue & stroma | ALL PAT | 0.65 | [0.609, 0.699] | 0.66 | 0.76 | 0.53 | <0.001 |
|  | ER+/HER2- | 0.65 | [0.556, 0.746] | 0.62 | 0.86 | 0.5 | 0.003 |
|  | ALL PAT (50) | 0.64 | [0.545, 0.729] | 0.62 | 0.82 | 0.4 | 0.004 |
|  | ER+/HER2- (50) | 0.58 | [0.517, 0.651] | 0.57 | 0.74 | 0.46 | 0.01 |
|  | DELTA | 0.60 | [0.483, 0.711] | 0.63 | 0.83 | 0.33 | 0.043 |
| tumor | ALL PAT | 0.65 | [0.608, 0.68] | 0.65 | 0.72 | 0.55 | <0.001 |
|  | ER+/HER2- | 0.65 | [0.562, 0.734] | 0.66 | 0.83 | 0.46 | 0.002 |
|  | ALL PAT (50) | 0.61 | [0.516, 0.712] | 0.57 | 0.82 | 0.38 | 0.014 |
|  | ER+/HER2- (50) | 0.55 | [0.49, 0.602] | 0.55 | 0.66 | 0.42 | 0.047 |
|  | DELTA | 0.62 | [0.516, 0.726] | 0.67 | 0.78 | 0.33 | 0.014 |
| stroma | ALL PAT | 0.78 | [0.731, 0.831] | 0.78 | 0.89 | 0.66 | <0.001 |
|  | ER+/HER2- | 0.74 | [0.684, 0.788] | 0.74 | 0.85 | 0.61 | <0.001 |
|  | ALL PAT (50) | 0.70 | [0.616, 0.786] | 0.70 | 0.88 | 0.55 | <0.001 |
|  | ER+/HER2- (50) | 0.64 | [0.569, 0.713] | 0.61 | 0.81 | 0.48 | 0.001 |
|  | DELTA | 0.69 | [0.524, 0.864] | 0.69 | 1.0 | 0.28 | 0.015 |

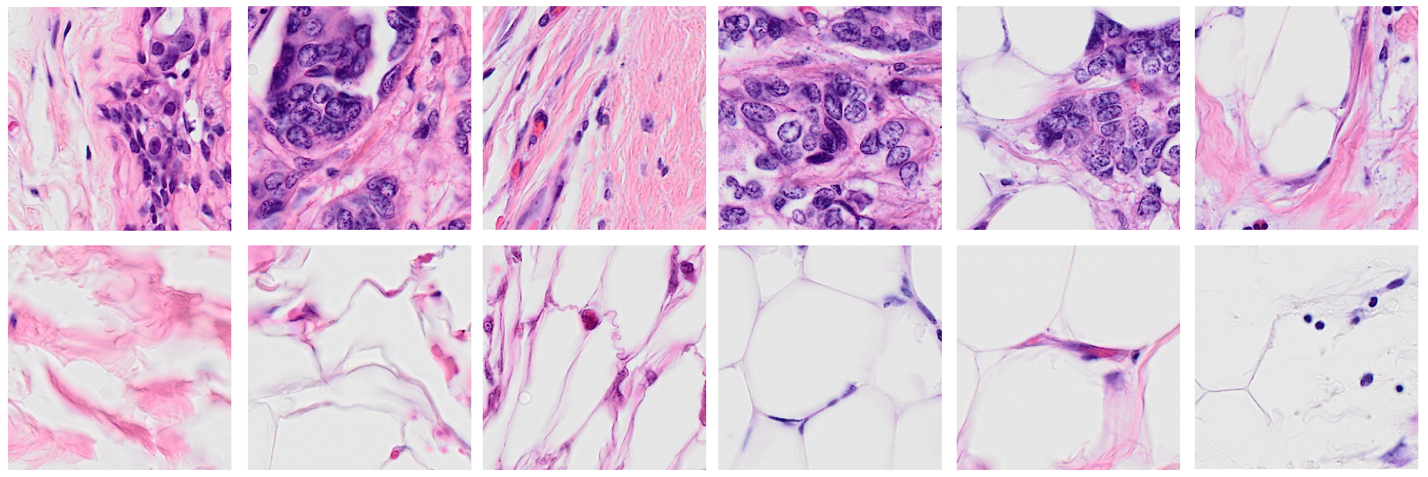

Figure S1: Examples of processed 380×380-pixel tiles. The top row shows tiles accepted as valid tissue samples using a pre-trained ResNet-50 tissue filter. The bottom row shows a set of rejected tiles. Tiles with mostly adipose tissue (2^nd^ row, 4^th^, 5^th^ and 6^th^ tiles) are removed from the dataset, but some residual adipocytes may remain in the filtered dataset (1^st^ row, 5^th^ and 6^th^ tiles).

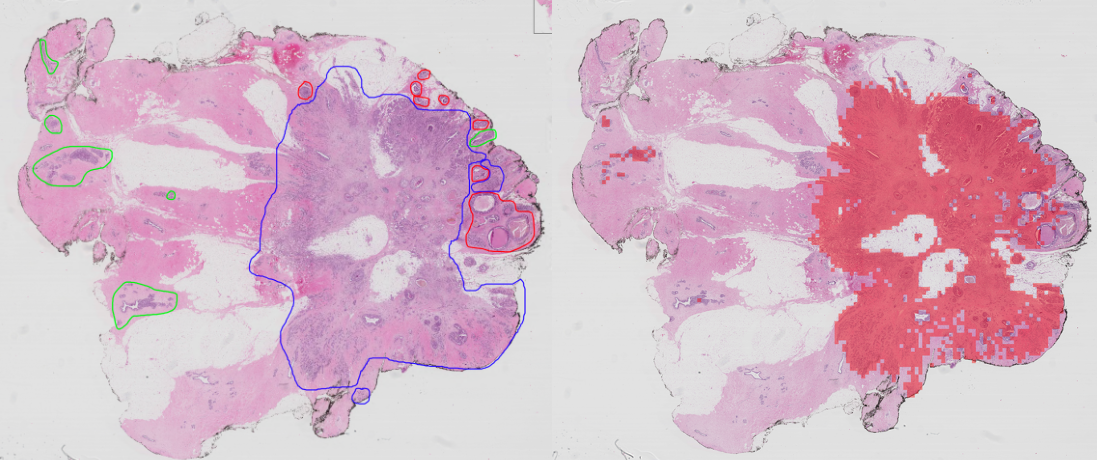

Figure S2: Example application of the CanvOI pan-cancer detector for classifying tiles as cancerous or non-cancerous. The left panel shows a whole-slide image from the BACH dataset, with tumor regions annotated by expert pathologists (Aresta, et al., 2019). The right panel displays the corresponding CanvOI predictions, with tiles classified as cancer overlaid in red.

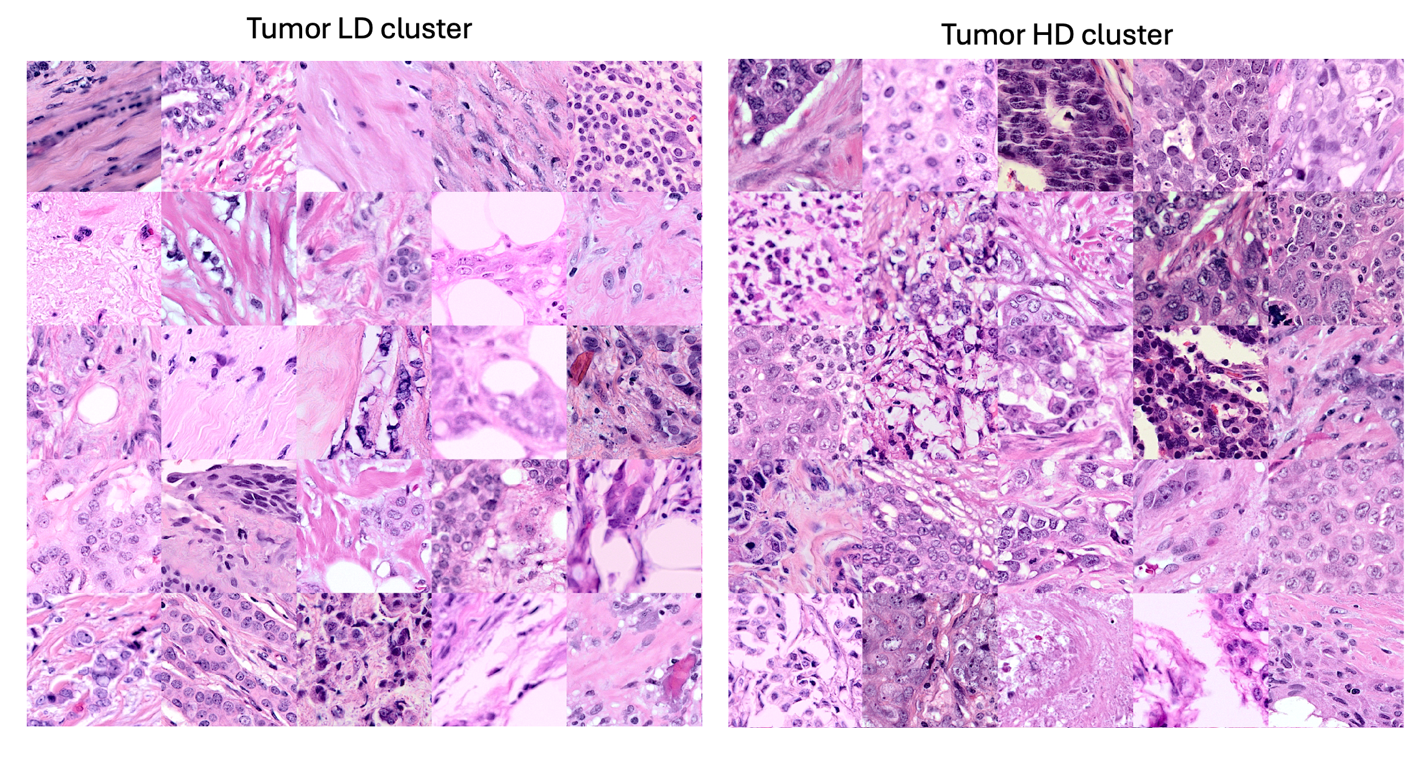

Figure S3: Examples of tumor tiles of size 380×380 pixels clustered using their representation vectors to approximately segregate low cellular density (tumor LD) and high cellular density (tumor HD) regions.

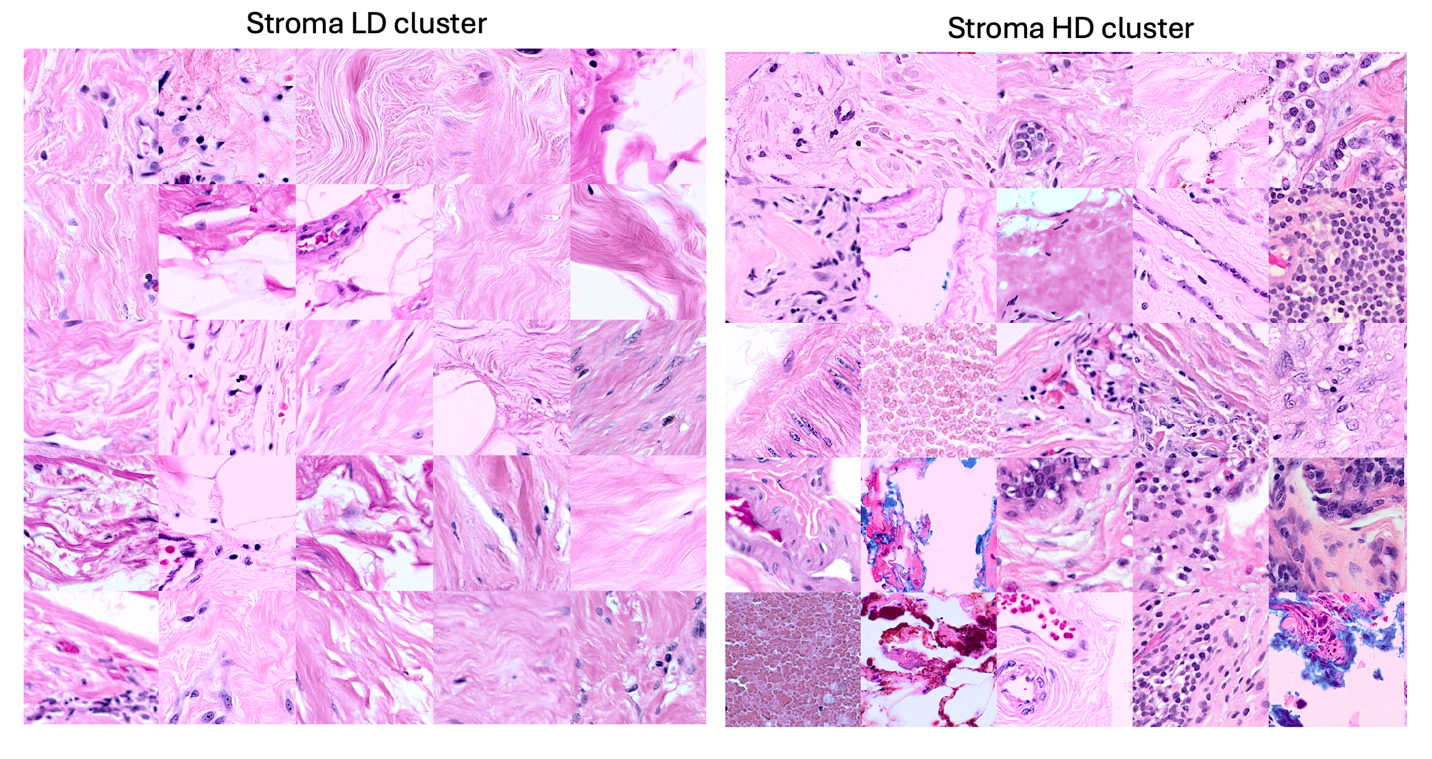

Figure S4: Examples of stromal tiles of size 380×380 pixels clustered using their representation vectors to approximately segregate low cellular density (stroma LD) and high cellular density (stroma HD) regions.
